# MicroRNA-transcriptome networks in whole blood and monocytes of women undergoing preterm labor

**DOI:** 10.1101/575944

**Authors:** Alison G. Paquette, Oksana Shynlova, Xiaogang Wu, Mark Kibschull, Kai Wang, Nathan D. Price, Stephen J Lye

## Abstract

Preterm birth is attributed to neonatal morbidity as well as cognitive and physiological challenges. We have previously identified significant differences in mRNA expression in whole blood and monocytes, as well as differences in miRNA concentration in blood plasma, extracellular vesicles (EV) and EV-depleted plasma in women undergoing spontaneous preterm labor (sPTL). The goal of this analysis was to identify differences in miRNA expression within whole blood (WB) and peripheral monocytes (PM) from the same population of women undergoing sPTL compared to nonlaboring controls matched by gestational age. We performed single end small RNA sequencing in whole blood and peripheral monocytes from women undergoing sPTL with active contractions(24-34 weeks of gestation, N=15) matched for gestational age to healthy pregnant non laboring controls (>37 weeks gestation, N=30) who later delivered at term as a part of the Ontario Birth Study (Toronto, Ontario CA). We identified significant differences in expression of 16 miRNAs in PMs and 9 miRNAs in WB in women undergoing sPTL. In PMs, these miRNAs were predicted targets of 541 genes, including 28 previously associated with sPTL. In WB, miRNAs were predicted to target 303 genes, including 9 previously associated with sPTL. These genes were involved in a variety of immune pathways, including interleukin 2 signaling. This study is the first to identify changes in miRNA expression in WB and PMs of women undergoing sPTL. Our results shed light on potential mechanisms by which miRNAs may play a role in mediating systemic inflammatory response in pregnant women that deliver prematurely.

## INTRODUCTION

Premature birth, defined as delivery before 37 weeks of gestation, occurs in 11.1% of pregnancies worldwide and is associated with neonatal morbidity and mortality[1]. Pregnancies characterized by pathological changes including placental insufficiency, subclinical infections, disruptions in maternal immune-tolerance to pregnancy, and decidual senescence often result in spontaneous preterm labor (sPTL). There is a paucity of research examining transcriptomic changes that occur during sPTL[2], which have the potential to transform our understanding of the molecular mechanisms underlying this heterogeneous syndrome [3]. Therefore, there is a crucial research need for robust and multidimensional characterization of transcriptomic changes relating to sPTL.

Throughout pregnancy, maternal blood circulates through the in-utero environment and responds to fetal cues. At the beginning of gestation, the number of monocytes in maternal blood drastically increase[4], which regulate placental invasion, angiogenesis, and tissue remodeling [5]. After infiltration into uterine tissues, monocytes differentiate into macrophages characterized by an immunosuppressive (M2) phenotype in normal pregnancies and an inflammatory (M1) state in complicated pregnancies[5]. Maternal immune cells play a crucial role in normal pregnancy maintenance and provide insight into changes that occur in pregnancies complicated by preterm birth.

Micro(mi)RNAs are a subtype of small, non-coding RNAs which are transcriptional regulators of gene expression in all human organs. MiRNAs are expressed within the placenta throughout pregnancy and are involved in fetal and maternal signaling. Placental miRNA expression profiles have been associated with preeclampsia [6] [7] and sPTL[8]. Like other organs, the placenta releases miRNAs into the circulation, and unique placenta derived miRNAs (from the C19MC and C14MC miRAN cluster) are detectable within maternal plasma[9–11]. Differential miRNA expression in plasma has been detected in pregnancies complicated by fetal growth restriction preeclampsia[13], and preterm birth[14, 15]. Circulating placenta enriched miRNAs are detectable in whole blood, and have been associated with fetal hypoxia[16]. MiRNAs may also play a role in monocyte differentiation throughout pregnancy, as there are differences in miRNA concentration in different monocyte subpopulations (CD16+ vs. CD16−)[17]. miRNAs have emerged as important transcriptional regulators and signaling molecules during pregnancy with the potential to play a role in the underlying molecular perturbations that occur during pregnancy-related complications such as sPTL.

Genome scale transcriptomic analysis of maternal blood provides a window into the changes that occur throughout pregnancy and during parturition. We have previously identified differences in mRNA expression between whole blood and peripheral monocytes[18], as well as differences in miRNA concentration in whole plasma, extracellular vesicles (EVs), and EV depleted plasma in women undergoing sPTL. The goal for this analysis was to identify changes in miRNA expression in maternal whole blood and monocytes in the same population (15 women undergoing sPTL and 30 pregnant women matched on gestational age not undergoing labor) and to integrate these results with our previous findings. We hypothesize that there are changes in miRNAs in whole blood and monocytes of women undergoing sPTL compared to controls, which are related to the role of these miRNAs in transcriptional regulation of labor-associated genes. Through this miRNA analysis and integration with next generation sequencing mRNA data from the same individuals, we obtain a better understanding of the transcriptional regulation that occurs in the context of PTL.

## MATERIALS AND METHODS

### Study Design

Participants were recruited within the Ontario Birth Study (OBS); a continuously enrolling prospective cohort at Mount Sinai Hospital (MSH) in Toronto Canada. Inclusion criteria for the OBS are: pregnancy diagnosed <17 weeks, maternal age >18, English speaking, signed informed consent, and intent to deliver and receive antenatal care at MSH. Exclusion criteria included non-viable neonate and/or inability to provide consent. Peripheral blood was collected from 15 patients who were in preterm labor, defined as cervical dilatation >4 cm and active uterine contractions, with no other accompanying pathology; who delivered prematurely between 24-34 weeks of gestation (Preterm labor or PTL). They were matched with 30 healthy asymptomatic pregnant women whose blood was taken at the same time point during routine clinical visits(gestational age 24-34 weeks), who later delivered at full term (TL). A full description of this subpopulation is described in other manuscripts [18] [19]. This study was approved by the Research Ethics Board of Mount Sinai Hospital, Toronto, Canada (#04-0024-E). All patients provided written consent as part of the OBS at Sinai Health System, Toronto, Canada.

### Specimen Collection

In PTL patients, peripheral blood samples were collected prospectively at the point of hospital admission, and in TL controls blood was collected during the regular antenatal visit. Blood was collected into both PAXgene tubes (Qiagen; Hilden, Germany) for whole blood isolation and EDTA blood collection tubes for monocyte separation. Monocytes were separated through the Monocyte RosetteSep cocktail (Stemcell Technologies, Vancouver CA), followed by high density gradient centrifugation to generate a highly purified monocyte fraction for subsequent mRNA isolation.

### Small RNA Sequencing and Quantification

Whole blood RNA was isolated using a PAXgene blood miRNA kit (Qiagen Hilden, Germany), and from peripheral monocytes using TRIzol LS reagent (Thermo Fisher, Waltham MA) following manufacturer’s instructions. RNA quality was determined using Experion analyzers (BioRad, Hercules CA) and sequenced at The Center for Applied Genomics at the SickKids Hospital, Toronto, Canada. Library preparation was performed following the New England Biolabs NEB Next multiplex small library preparation protocol, with 400 ng of total RNA as the input. The 3’ adapter was ligated to the small RNA, followed by reverse transcriptase (RT) primer hybridization to the 3’ adapter and then the 5’ adapter was ligated to the opposite end. Libraries were generated using first stranded synthesis then enriched by PCR. Quality and size was determined using the Bioanalyzer 2100 DNA High Sensitivity chip (Agilent Technologies, Palo Alto CA), and libraries were quantified by qPCR using the Kapa Library Quantification Illumina/ABI Prism Kit protocol (KAPA Biosystems, Wilmington MA) and sequenced on Illumina HiSeq 2500 (Illumina, San Diego CA).

Sequencing data was preprocessed using the RNA analysis pipeline sRNAnalyzer ([20], http://srnanalyzer.systemsbiology.net/). In the data preprocessing, adaptor sequences were trimmed and low-quality sequences were removed. Processed sequences were aligned to all human miRNAs (miRBase Release 21,[21]) with no mismatches. We removed miRNAs with 0 read counts in >50% of samples or a mean read count <20 producing a final dataset of 417 miRNAs in whole blood and 274 miRNAs in peripheral monocytes, CPM (Count Per Million) values were calculated using RNA sequencing analysis software “edgeR”[22], and log2 transformed. Data from this analysis is publicly available within the Gene Expression Omnibus (GEO) as GSE108876 and GSE108877.

### Statistical Analyses

Differentially expressed miRNAs were identified using edgeR [22]. miRNAs were considering significantly associated with PTL if they exhibited a Benjamini Hochberg (BH) adjusted *q* value of <0.05, and a log2 fold change of >1. In order to mitigate batch effects, we eliminated differentially expressed miRNAs which were correlated with order they were loaded onto the array (N=3 miRNAs in whole blood and monocytes), identified using spearman correlation tests with a statistical cutoff of FDR adjusted *q* value<0.05. We compared expression of miRNA in whole blood and peripheral monocytes to previously published data from the same patient population, including miRNA data from matched plasma and EVs(GSE106224)[19], as well as mRNA expression from whole blood and peripheral monocytes (GSE96097) [18]. Putative miRNA targets were detected using the quantitative model TargetScan (V. 7.0, targetscan.org), which characterizes canonical targeting of miRNAs based on 14 features, and has the best predictive performance compared to comparable tools[23]. MiRNA-target gene relationships were identified in whole blood and peripheral monocytes by examining correlations between each miRNA and its proposed target genes. For each miRNA, we used only mRNA targets with an absolute value of “context++” score (a metric of miRNA target prediction accuracy used by TargetScan) which was higher than the median, ensuring only highest quality relationships were validated. miRNA and target relationships were considered statistically significant using a Spearman correlation coefficient <−0.3 and a *p* <0.05. These mRNAs were then matched to the list of Differentially Expressed Genes (DEGs) identified in whole blood and monocytes from the same individuals [18].

Gene set enrichment analysis was performed for mRNA targets using two-sided hypergeometric tests conducted on Gene Ontology (GO) Biological Process Gene sets using GO gene set visualization application “ClueGO” within the cytoscape environment [24]. GO gene sets with more than 5 genes were included in the analysis, and were considered significant with a Benjamini Hochberg adjusted *q* <0.05. These GO gene sets were clustered into ClueGO groups based on the similarity in the number of genes in the gene sets calculated by a Kappa score within ClueGO. Data was analyzed and visualized in R (Version 3.3.1) and Cytoscape (Version 3.6.0) [24].

## RESULTS

### Population Characteristics

Demographic characteristics of women undergoing sPTL (defined as active uterine contractions with cervical dilation followed by delivery between 24-34 weeks) were compared to healthy women not in labor that went on to deliver at full term (**Table 1**). There were no significant differences in delivery method, ethnicity, or fetal sex between sPTL cases and controls (*p*>0.05, Fisher’s exact test). Women who delivered prematurely were more likely to be between 18 and 30 years old vs. older than 30, although this relationship was borderline statistically significant (*p*=0.06).

**Table 1:**
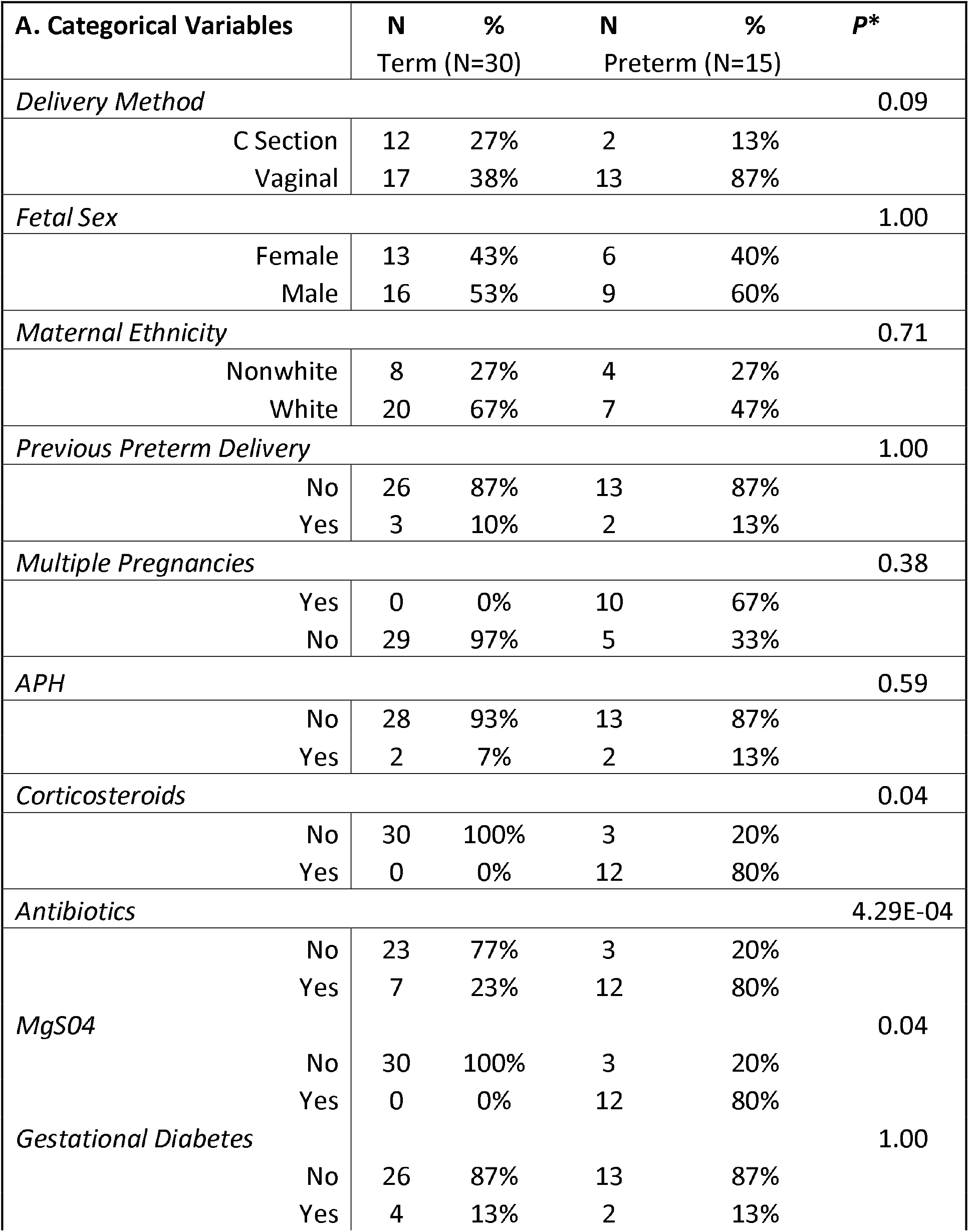
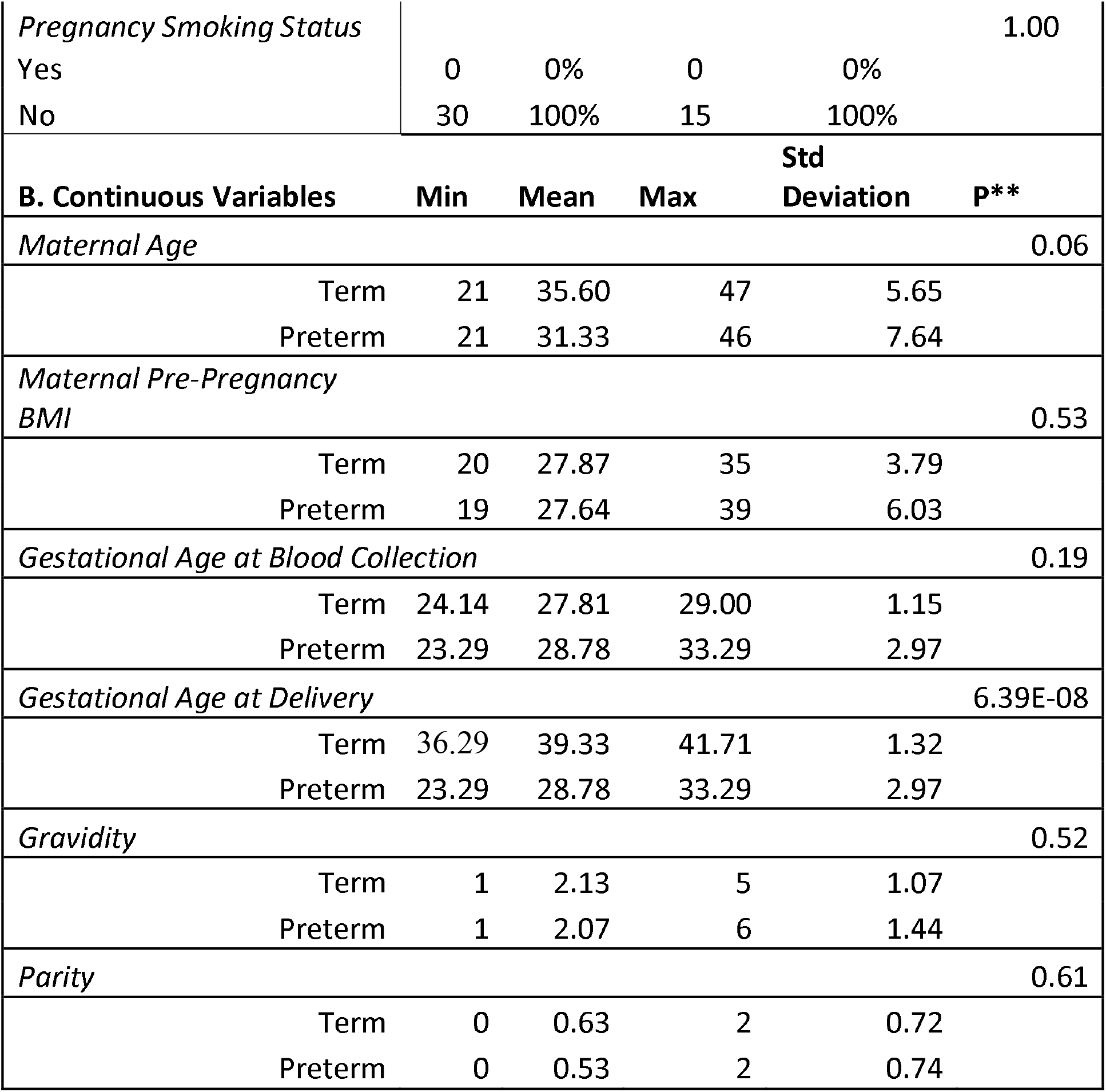
Participant Characteristics

### Differences in miRNA expression in Whole Blood and Monocytes

We identified miRNAs with significantly different concentrations in peripheral monocytes and whole blood of women undergoing PTL using generalized linear models within edgeR [22]. In monocytes, 11 miRNAs were higher and 5 miRNAs were lower in women undergoing PTL compared to women who delivered at term (***Figure 1A-B, supplemental table 1***). In whole blood, 6 miRNAs were higher and 3 miRNAs were lower in women undergoing PTL compared to controls (***Figure 1 C-D, supplemental table 1***). Two of these miRNAs (mIR-495-3p and miR-381-3p) were from the C14MC MC miRNA cluster; a region of chromosome 14 which encodes miRNAs which are substantially enriched in placental tissue. Overall, there was a higher number of significant miRNAs and larger concentration changes in relation to sPTL in monocytes compared to whole blood.

**Figure 1:**
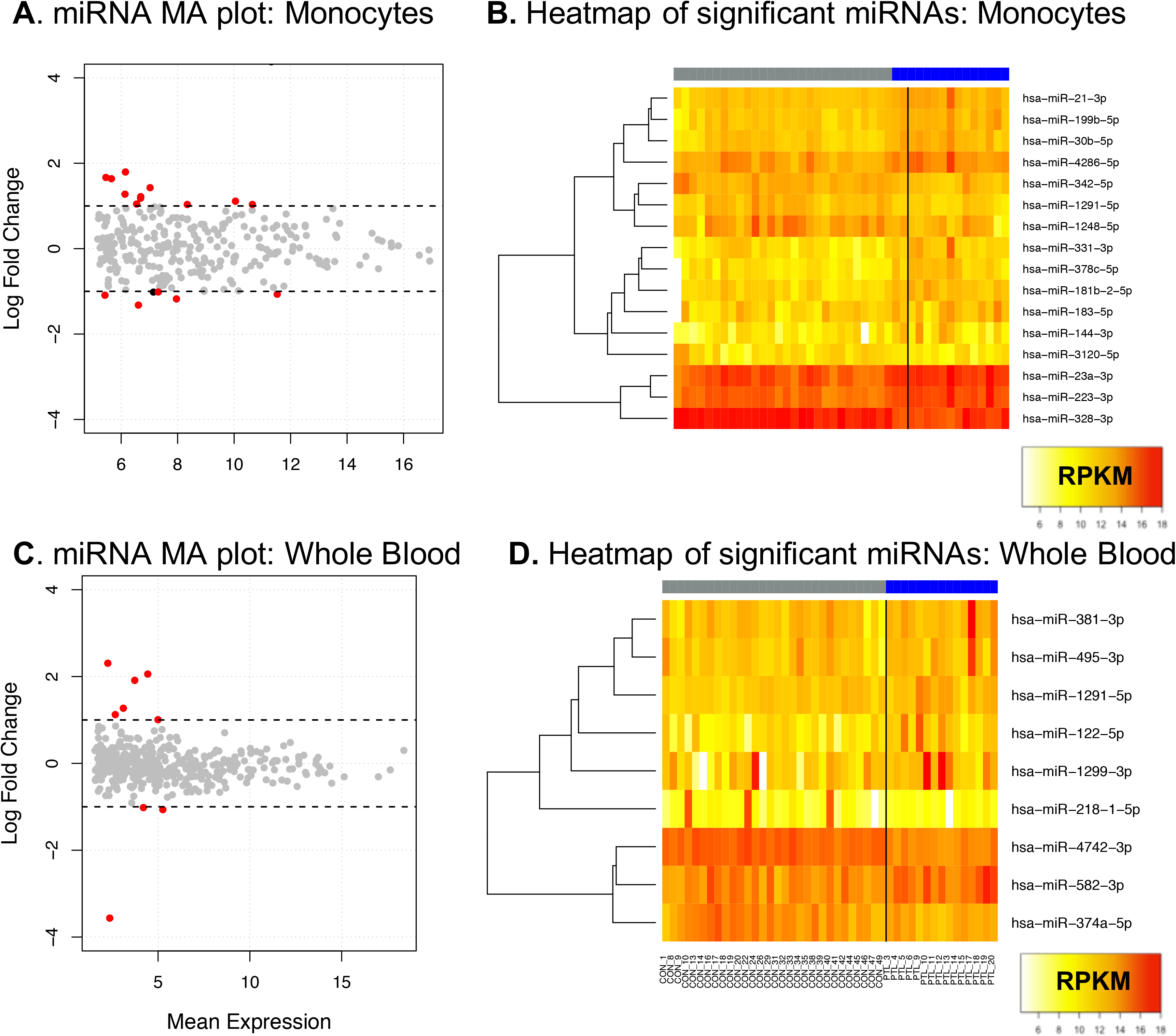
MA plot of log2 fold changes vs. the mean of normalized counts in the regularized logarithmic distribution of RNA sequencing data in (**A**) monocytes, and (**C**) whole blood. mRNAs significantly different in linear models after adjustment for multiple comparisons are highlighted in red, and lines indicate log2 fold changes >1. Heatmap of the 16 differentially expressed genes identified in monocytes (**B**) or the 9 differentially expressed genes identified in whole blood (**D**), with blue indicating lower expression and red indicating higher expression. miRNA expression of women who underwent term labor is on the left and highlighted in grey, and miRNA expression of women in sPTL is highlighted in blue on the right.

MiR-1291-5p expression was significantly increased in whole blood of women undergoing sPTL(Log fold change 1.12, FDR adjusted *q*=2.22×10^−4^), but decreased in monocytes of women undergoing sPTL (Log fold change −1.32, FDR adjusted *q*=2.45×10^4^). We observed a significant negative correlation between expression of this miRNA in whole blood and in monocytes (*p*=0.02, ρ =−0.02, Spearman correlations, **Supplemental Figure 1**). This higher level in whole blood suggests that other sub-populations of blood cells (such as erythrocytes, platelets, lymphocytes or granulocytes) may have the opposite expression profile compared to monocytes and also contribute to the miRNA pool in women undergoing sPTL.

### Comparison of expression differences across blood compartments

In our prior analysis of plasma from same patients from the OBS cohort, we identified significant differences in concentration of 132 miRNAs in whole plasma, EVs and EV depleted plasma [19]. Five of these miRNAs were also significantly different in monocytes, and three miRNAs were significantly different in whole blood (**supplemental table 2**). We examined correlations of these 8 miRNAs that were significantly upregulated in plasma samples as well as in whole blood and monocytes from the same patients (**supplemental figure 2**, N=44 samples with plasma and whole blood RNAseq data, N=42 samples with monocyte and whole blood sRNA sequencing data, and N=14 samples with RNAseq data in EVs and monocytes/whole blood).

Two miRNAs (miR-183-5p, and miR-331-3p) were increased and one miRNA (miR-328-3p) was decreased in both plasma and monocytes of women undergoing sPTL compared to TL controls. The concentration/expression of these miRNAs was not significantly correlated between whole plasma and monocytes (*p*>0.05, Spearman correlations). MiR-181b-5p was increased in monocytes and decreased in the plasma of women undergoing sPTL, and the expression/concentration of these miRNAs was inversely correlated in plasma and monocytes (ρ= −0.28, *p*=0.07). MiR-378c-5p was increased in both monocytes and in extracellular vesicles of women undergoing sPTL. We observed significant positive correlations (*p*<0.05) between whole blood miRNA levels and the levels in plasma and EVs in the 3 miRNAs significantly associated with sPTL in the previous studies: (1) MiR-374a-5p was decreased in both plasma and whole blood of women undergoing sPTL, (2) miR-381-3p was decreased in plasma but increased in whole blood of women undergoing sPTL, and (3) miR495-5p was decreased in the plasma and extracellular vesicles but increased in the whole blood of women undergoing sPTL compared to controls. This indicates that there is some overlap in signal related to sPTL that is detectable across different components of the blood within the same individuals.

### Confirmation of target mRNAs of miRNAs associated with sPTL

We identified the putative mRNA targets of the differentially expressed miRNAs using the TargetScan database. TargetScan incorporates 14 features to predict miRNA-mRNA interactions and has shown to have the highest predictive value compared to similar tools [23]. From the TargetScan database, we included only the top 50% of predicted mRNA based on the “context ++” score (a metric of accuracy used by the database). We then compared mRNA expression of these predicted targets to the miRNA concentration within matched samples (GSE96097,[18]). We constrained miRNA-mRNA predicted interactions to only those which had a Spearman correlation coefficent ρ < −0.3 and a *p* <0.05, which represented 7.29% of the predicted genes in monocytes and 4.26% in whole blood (**Supplemental table 3, Supplemental Figure 4**). In monocytes, 541 genes were associated with 16 miRNAs, and in whole blood 303 genes were associated with 7 miRNAs. Based on this criteria, there were 2 miRNAs in whole blood with no confirmed mRNA targets.

In previous transcriptomic analyses from the same patients, we identified 262 genes in monocytes and 181 genes in whole blood which were associated with sPTL[18]. Here, we compared the changes in mRNA expression related to sPTL and the confirmed miRNA target genes we have identified in this current study (**Figure 2**). In whole blood, 9 of the 181 DEGs were confirmed miRNA targets of 3 differentially expressed miRNAs in whole blood (Shown in **Figure 2A**). Among these 3 miRNAs in whole blood, miR-4742-3p emerged as the strongest regulator, since it was negatively associated with 5 genes which were positively associated with sPTL, including *SPH, CD177, CYP1B1, ELOVL7* and *GRB10*. In monocytes, 28 of these 262 genes were confirmed mRNA targets of 7 differentially expressed miRNAs. In monocytes, MiR-1291-5p emerged as the strongest regulator, and was negatively associated with 11 of the 28 genes which are positively associated with sPTL, including *IL1B, IL1R1* and *CD177* (**Figure 2B**). In both whole blood and monocytes, the direction of the associations with PTL is congruent with the directionality in mRNAs (i.e., if a miRNA is positively associated with PTL, the miRNA is negatively correlated with mRNAs that are negatively associated with PTL). We also examined mRNAs which were in the same genomic region as the miRNAs in our study, but found that none of the associated mRNAs were statistically significantly associated with sPTL in monocytes and whole blood leukocytes. This integrated analysis suggests that the miRNAs identified here may play a role in the transcriptional regulation of genes associated with sPTL in monocytes and whole blood leukocytes.

**Figure 2:**
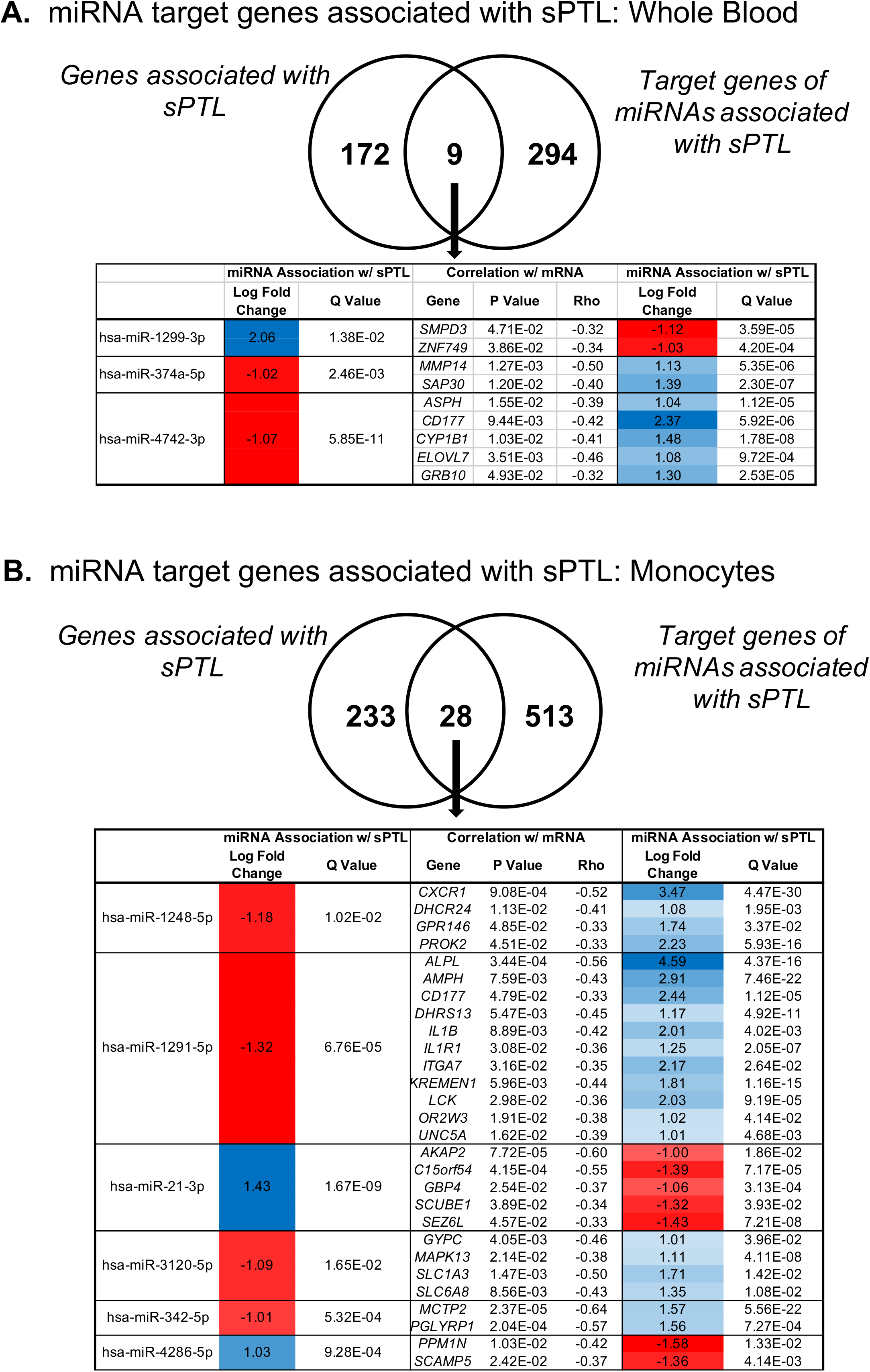
**A.** Venn diagram in whole blood (**A**) and monocytes (**B**) of genes previously associated with sPTL and gene targets of the differentially expressed miRNAs identified in this study. In the overlapping genes, we show the log fold change and P values of the miRNAs (from supplemental table 1), correlation with mRNA concentration, and log fold change and correlation with sPTL in the previous study).

### miRNA target mRNA networks in sPTL

We performed enrichment analysis to identify gene sets which were overrepresented by the confirmed mRNA targets of the differentially expressed miRNAs identified in whole blood and monocytes. Using CytoScape application “ClueGO”, we identified significantly enriched GO gene sets which were grouped together based on intersecting common genes, which are shown in **Figure 3, and in supplemental table 4**. In whole blood, 59 different GO gene sets were significantly enriched for the 303 genes which were mRNA targets for the 9 miRNAs associated with sPTL in whole blood. These gene sets were grouped into 19 ClueGO groups, including 8 distinct groups (i.e. only had 1 GO gene set), and one ClueGO group which contained 22 GO gene sets including the 5 most significant gene sets: “positive regulation of cytokine production”, “positive regulation of adaptive immune response”, “interleukin-2 production”, “positive regulation of adaptive immune response based on somatic recombination of immune receptors built from immunoglobulin superfamily domains”, and” T cell receptor signaling pathway”. This diversity of GO gene sets and strong signal indicates that these miRNAs associated with sPTL are involved in a wide variety of biological pathways in whole blood; particularly immune pathways.

**Figure 3:**
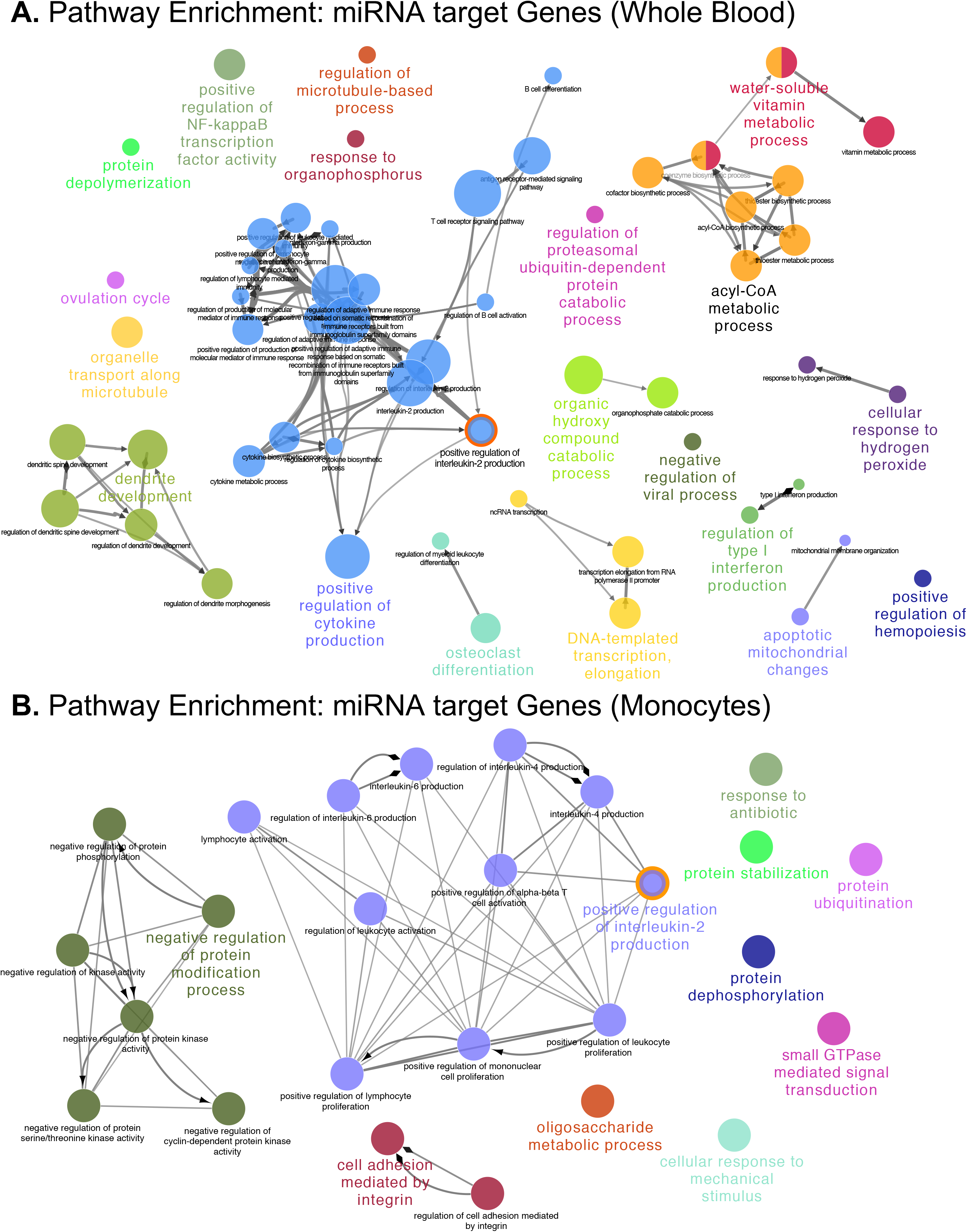
Biomolecular network of miRNA target genes in (**A**) monocytes and (**B**) whole blood. Enrichment analysis was performed on gene ontology (GO) gene sets and significant gene sets are shown (Benjamini Hoch adjusted q<0.05). Network node layout is based on similarity between genes within GO gene sets, where nodes are colored based on multiple occurrences within different go categories. GO terms which were significant in both whole blood and monocytes have a red outer circle.

In monocytes, 30 different gene sets were significantly enriched for the 541 genes which were targets for the 16 miRNAs associated with sPTL. These gene sets were clustered into 11 groups, including 8 gene sets which were distinct (i.e only had 1 GO term), and 1 gene set (Group 10), which had 13 different terms. The top 5 GO terms based on q value were: “small GTPase mediated signal transduction”,” negative regulation of protein modification process”, “positive regulation of interleukin 2 production”, “positive regulation of mononuclear cell proliferation”, and “negative regulation of protein kinase activity”. These terms are reflective of changes associated with monocyte proliferation and immune function, indicating that miRNAs may play a role in regulation of these pathways in monocytes.

The Gene Ontology gene set “positive regulation of interlukin-2 production” was significantly enriched in both whole blood and monocytes. This gene set contains 31 genes, and the differentially expressed miRNAs in both whole blood and monocytes were predicted negative regulators of 5 unique genes within this gene set, each representing 17.86% of the total genes in this pathway. In whole blood, 4 miRNAs were identified as regulators of the 5 genes (*CCR2, CD4, MALT1, PDE4B*, and *CD28*). In monocytes, 3 miRNAs were identified as negative regulators of the 5 genes (*SASH3, CD83, IL1B, TNFSF4, and CD28*) involved in interleukin signaling (**Table 2**). In this gene set, *IL1B* was a target of miR-1291-5p, and *IL1B* expression was significantly increased in women undergoing sPTL (FDR adjusted *q*=4.02×10−3, logFC=2.01). Through this analysis, we have identified that miRNA expression within whole blood and peripheral monocytes significantly alters genes involved in production of interleukin 2.

**Table 2:** mRNA targets of differentially expressed miRNAs involved in Positive regulation of Interleukin-2 signaling

| miRNA | Gene targets in GO:0032743 |
| --- | --- |
| *Monocytes* |  |
| hsa-miR-1248-5p | *SASH3, CD83* |
| hsa-miR-1291-5p | *IL1B* |
| hsa-miR-378c-5p | *CD28, TNFSF4* |
| *Whole Blood* |  |
| hsa-miR-4742-3p | *CCR2* |
| hsa-miR-1299-3p | *CD4, MALT1* |
| hsa-miR-582-3p | *PDE4B* |
| hsa-miR-1291-5p | *CD28* |

## DISCUSSION

This study is the first comprehensive miRNA profiling of women undergoing sPTL using small RNA sequencing within both whole blood and peripheral monocytes. Our key findings included (A) identification of differentially expressed miRNAs in whole blood and peripheral monocytes, which (B) were negative regulators of genes associated with sPTL, which we identified using previously generated RNA sequencing data on whole blood and monocytes from the same individuals and (C) identification of unique and shared gene sets enriched for mRNA targets of these differential miRNAs in monocytes and whole blood of women undergoing sPTL.

Overall, we identified a greater number of miRNAs with concentration changes, and more genes related to miRNAs associated with sPTL. Additionally, we observed no congruence in the differential miRNAs identified in monocytes and whole blood. Maternal blood includes three major subsets of immune cells; monocytes, lymphocytes, and granulocytes, which circulate through gestational tissues including the myometrium, decidua and placenta, and are exposed and respond to signals from these tissues. RNA sequencing data derived from monocytes exhibits less cellular heterogeneity and thus likely has a cleaner signal. We suggest that the signaling we observe in monocytes may be related to their functional role throughout pregnancy and parturition [4]. Altogether, our work suggests that a stronger signal related to sPTL can be obtained from monocytes compared to whole blood.

We observed a number of miRNAs which were differentially expressed in in whole blood and monocytes of women undergoing sPTL which also exhibited significant concentration differences previously identified in whole plasma, EV-depleted plasma and EVs in the same population of individuals [19]. We identified fewer differential miRNAs in immune cells compared to the plasma component, but overlap in the differentially expressed miRNAs, as well as concurrence in the directionality of associations with sPTL. This suggests functional signaling that occurs within blood in context of sPTL. In independent populations, increased expression of miR-223 has been identified in plasma of women at 20 weeks who went on to deliver prematurely[14]. Similarly, we observed increased levels of this miRNA in whole blood of women who delivered prematurely compared to TL controls, which supports the potential of this miRNA as having a functional role in PTL.

In whole blood, 2 of the miRNAs (miR-495-3p and miR-381-3p) with concentration differences in women undergoing sPTL were part of the C14MC region; a group of imprinted miRNAs located on chromosome 14q32 which is highly produced by the placenta and enters maternal circulation [25]. In plasma samples from the same patients, these miRNA concentrations were negatively associated with sPTL[19], and changes in average expression of C14MC miRNAs in plasma have also been associated with preterm labor[15]. On average, C14MC miRNAs have higher expression and lower variance in the fetal compartment and the placenta compared to the maternal plasma, thus this signal may be reflecting changes in this compartment[26]. We hypothesize that the changes observed within the whole blood and monocyte fraction of leukocytes are reflective of functional transcriptional changes that occur within the blood or they may be reflective of concentration changes related to miRNA signals from the developing fetus.

We identified mRNAs which were transcriptionally regulated by differentially expressed miRNAs using predicted targets from TargetScan, which were filtered using matched mRNA data derived through RNA sequencing based on negative correlations with miRNA[18]. Of the targets identified by TargetScan, on average only 4.26% of these predicted targets passed our filter whole blood and 7.29% in monocytes. These target prediction algorithms are only designed to reveal gene expression regulation potential, and expression can also be altered by post transcriptional modifications including transcriptional regulation, histone acetylation and DNA methylation [27], and are not cell type specific [23]. This analysis was designed to collect miRNA and mRNA from the same patient, which allowed us to perform correlations and obtain a far more accurate understanding of miRNA-mediated regulation of gene expression than by relying solely on miRNA prediction algorithms alone.

We used our curated, confirmed miRNA target lists generated to perform an enrichment analysis of gene sets related to our differentially expressed miRNAs. In whole blood, we identified immune related gene sets. This may be reflective of the inflammatory and immune response that occurs as a breakdown in maternal fetal tolerance or inflammation related to preterm labor[28]. The target genes of the miRNAs differentially expressed within monocytes were enriched for a total of 19 GO gene sets including those related to interleukin production and activation of the immune system, such as “regulation of interleukin production (Interleukins 2, 4 and 6), as well as T Cell, lymphocyte, and monocyte proliferation. The monocyte population expands during pregnancy and differentiates into a pro-inflammatory phenotype during labor [30]. Our work indicates that part of this expansion in monocytes may be transcriptionally regulated by a subset of miRNAs in monocytes. More work is required to understand if these miRNAs induce monocyte expansion and proliferation *in vitro*, which is beyond the scope of this analysis.

In monocytes, miR-1291-5p emerged as a master miRNA regulator of the highest number of genes associated with sPTL, including *IL1B* and its receptor *IL1R1*, which have previously been associated with sPTL. The positive regulation of interleukin 2 gene set was significantly enriched for miRNA target genes in both whole blood and monocytes, which contains *IL1B*. This gene set is involved in immune response and activation of inflammatory cytokines [31]. Inflammatory cytokines increase at the end of pregnancy as monocytes shift into an inflammatory M1 phenotype [30], and are implicated in sPTL[32]. Our work suggests that this may be in part modulated through changes in miRNAs expression in whole blood and monocytes.

We are limited in our ability to perform extensive characterization, stratification/clustering or any predictive analyses of these miRNAs due to inadequate statistical power resulting from a small sample size. Additionally, we cannot separate miRNA signals related to PTL from those related to the labor process (uterine contractions) or to the betamethasone treatment that women delivering prematurely received prior to labor, as previously discussed [18]. Due this these limitations, this work was not designed to interrogate miRNAs as a potential biomarker for sPTL. Further functional analyses are required, including functional validation of miRNA/mRNA targets, as well as the functionality of these miRNAs on key tissues involved in pregnancy (i.e. placenta and myometrium). Additionally, our findings will need to be validated in independent cohort of pregnant women. This work instead focuses on potential mechanisms by which miRNAs may play a role in mediating systemic inflammatory response in pregnant women that deliver prematurely, serves as an important foundation to subsequent analyses, and highlights the importance of sample source when performing blood related assessments.

Our analysis stands out among related studies because it is the first to generate high throughput RNA sequencing data of both miRNA and mRNA from the same patient source. We have carefully selected a reliable methodology to identify potential miRNA targets based on experimental data [23], and have used novel alignment tools to obtain comprehensive coverage of miRNA expression [20]. This robust, multi-dimensional data has allowed us to confidently build mRNA-miRNA regulation networks related to sPTL in a way that has not previously been done, which provides additional insight into mechanisms of transcriptional regulation of human labor. This type of integrative analysis has been suggested to overcome many challenges in performing pregnancy research [3]. Our analysis has highlighted miRNA mediated transcriptional regulatory networks of sPTL-associated genes in monocytes and whole blood, which are involved in important biological pathways, including interleukin signaling. Further work is needed to validate these data in independent populations and elucidate the miRNA mechanisms of signaling in whole blood and in a subpopulation of peripheral monocytes.

## ACKNOWLEDGEMENTS

The authors would like to thank the women who participated in this study as well as the staff at Mount Sinai Hospital, in particular Drs Robin Thurman and Kellie Murphy. We would like to thank members of the Price lab, particularly Ben Heavner and Rory Donovan for their help with the data management and members of Lye lab, particularly Mrs Elzbieta Matysiak-Zablocki and Mrs Anna Dorogin for their help with blood processing. We would like to acknowledge Drs Lee Hood, Stephen Matthews, Sam Mesiano, Leslie Myatt, Craig Pennell and Yoel Sadovsky on behalf of the GAPPS “Systems Biology of Preterm Birth” pilot study, as well as Mary Brunkow for her coordination of this study. We would also like to thank Beverly Anne Bessie and Theresa Fitzgerald for their help with the coordination of this manuscript. This work is supported by the Global Alliance to Prevent Prematurity and Stillbirth (GAPPS), Grant Title “Systems Biology of Preterm Birth: A Pilot Study”, as well as the Eunice Kennedy Shriver National Institute of Child Health & Human Development of the National Institutes of Health: Grant R01HD091527 (to N.P), and K99HD096112 (to A.P).

---

**Supplemental Figure 1:** miR-1291-5p expression in whole blood and monocytes. miRNA expression values for women who delivered at term are highlighted in orange, and values for women who delivered prematurely are in blue.

**Supplemental Figure 2:** Correlations in miRNA expression which were identified as significant in (A) monocytes and plasma (B) monocytes and exosomes (C) whole blood and plasma/exosomes. Blue dots indicate correlations involving monocytes, and red dots indicate correlations involving whole blood. Orange dots reflect correlation between whole blood and exosomes. Circled dots are expression values involving exosomes. Correlations were evaluated using spearman correlations, with significance defined as *p*<0.05. All correlations; including those which are insignificant are shown here.

**Supplemental Figure 3:** Number of miRNA targets identified in (A) whole blood and (B) Monocytes. Blue lines indicates the number of targetscan candidates which had higher than the median context ++ scores, which is a measure of prediction accuracy to bind to target miRNA. Green bars indicate mRNAs with matched expression data in GSE96097. Yellow bars indicate samples whose miRNA expression in whole blood and/or monocytes was significantly correlated with the mRNA expression (defined as ρ <−0.3, *p*<0.05, spearman correlation coefficients) in matched samples.

