## Supplementary figures and images for "MicroRNA-transcriptome networks in whole blood and monocytes of women undergoing preterm labor"

### Supplemental Figure 1

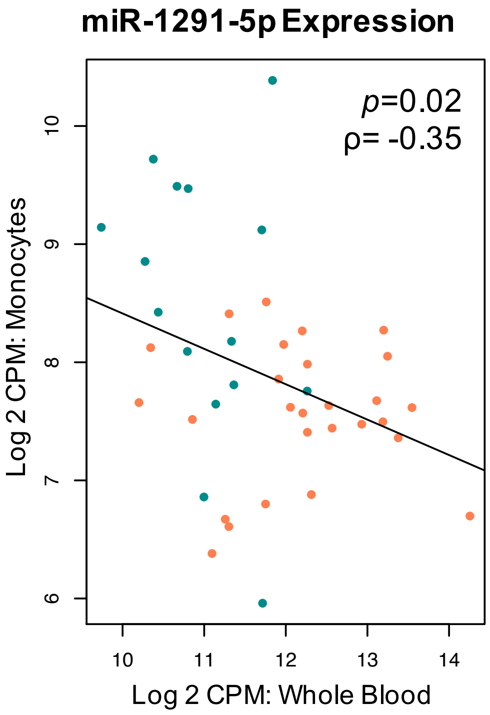

### Supplemental Figure 2

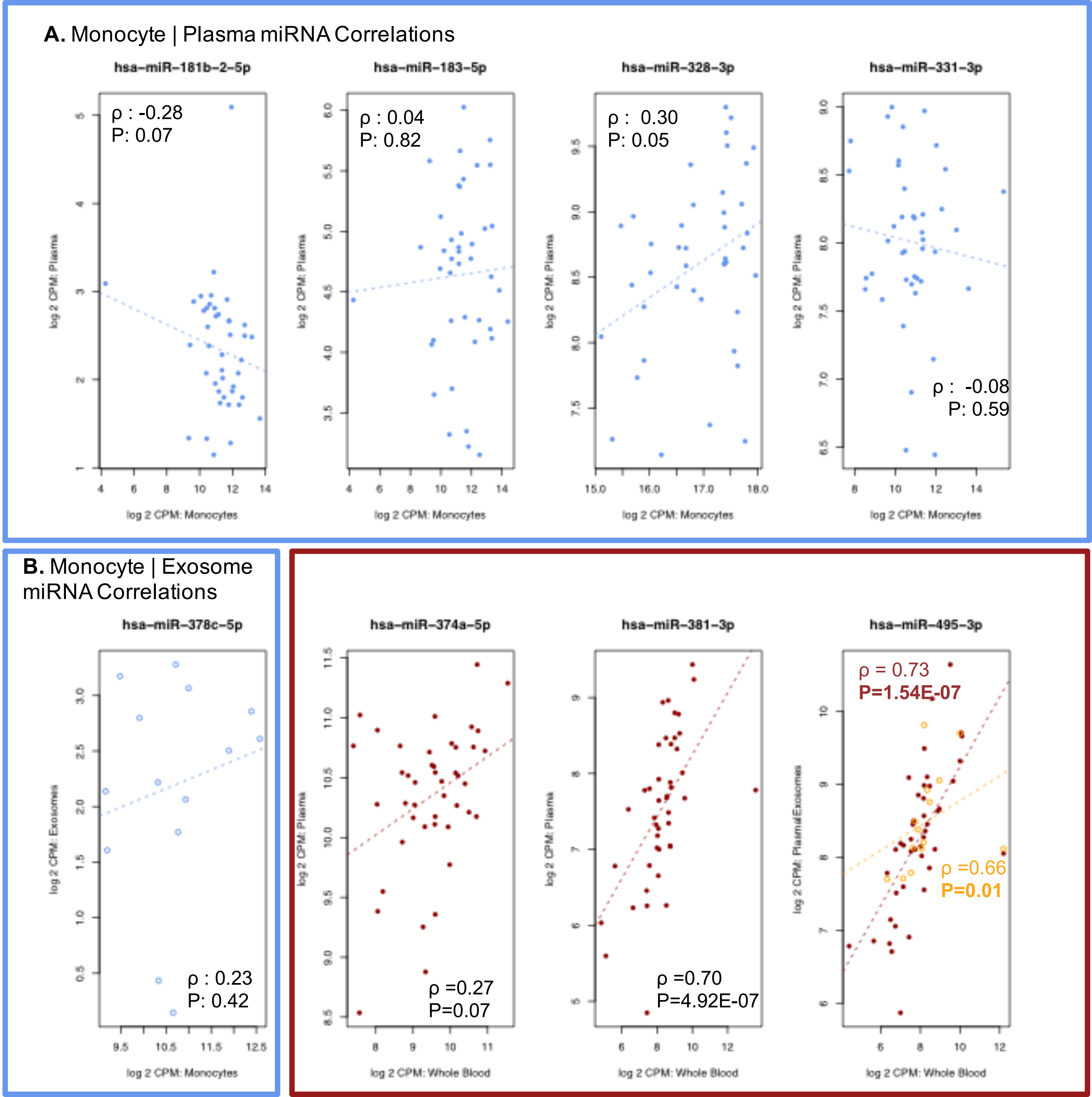
