## Supplemental Figure 3 for "MicroRNA-transcriptome networks in whole blood and monocytes of women undergoing preterm labor"

**A.** Confirmation of predicted mRNA targets of miRNAs in Whole Blood

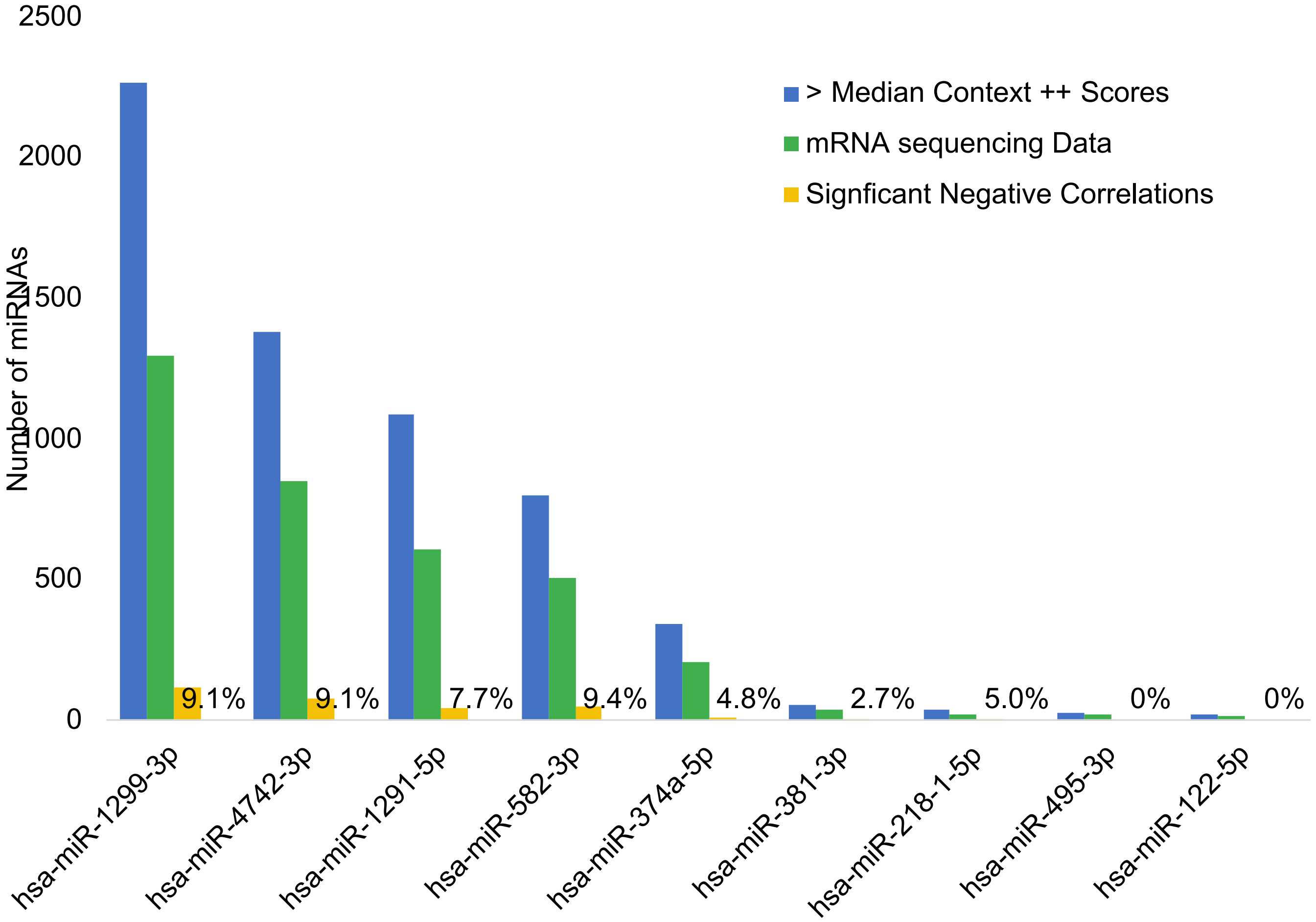


**B.** Confirmation of predicted mRNA targets of miRNAs in Monocytes

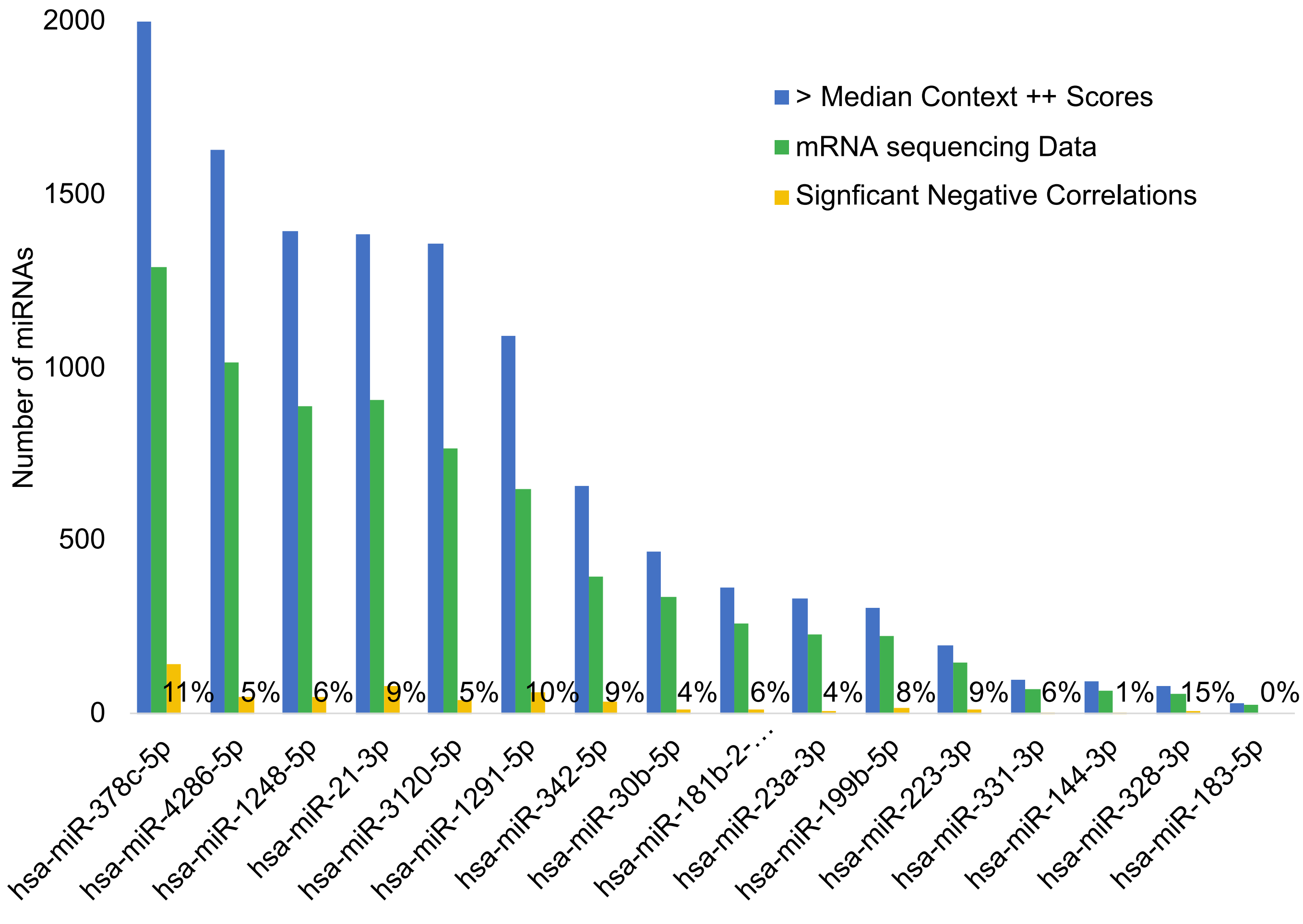
